# Overexpression of the *WAPO-A1* gene increases the number of spikelets per spike in bread wheat

**DOI:** 10.1101/2022.04.01.486512

**Authors:** Lukas M. Wittern, Jose M. Barrero, William D. Bovill, Klara L. Verbyla, Trijntje Hughes, Steve M. Swain, Gareth Steed, Alex A.R. Webb, Keith Gardner, Andy Greenland, John Jacobs, Claus Frohberg, Ralf-Christian Schmidt, Colin Cavanagh, Antje Rohde, Mark Davey, Matthew A. Hannah

**Affiliations:** Department of Plant Sciences, University of Cambridge, CB2 3EA, UK; BASF, BBCC - Innovation Center Gent, Technologiepark 101, 9052 Ghent, Belgium; National Institute of Agricultural Botany (NIAB), Huntingdon Road, Cambridge, CB3 0LE, UK; Commonwealth Scientific and Industrial Research Organisation, Agriculture and Food, Black Mountain Science and Innovation Park, Canberra, ACT 2601, Australia; BASF Australia Ltd., 28 Freshwater Place, 3006 Melbourne, Australia

**Keywords:** *WAPO1*, spikelets per spike (SPS), spike architecture, Multiparent Advanced Generation Intercross (MAGIC), wheat transgenics

## Abstract

Two homoeologous QTLs for number of spikelets per spike (SPS) were mapped on chromosomes 7AL and 7BL using two wheat MAGIC populations. Sets of lines contrasting for the QTL on 7AL were developed which allowed for the validation and fine mapping of the 7AL QTL and for the identification of a previously described candidate gene, *WHEAT ORTHOLOG OF APO1* (*WAPO1*). Using transgenic overexpression in both a low and a high SPS line, we provide a functional validation for the role of this gene in determining SPS also in hexaploid wheat. We show that the expression levels of this gene positively correlate with SPS in multiple MAGIC founder lines under field conditions as well as in transgenic lines grown in the greenhouse. This work highlights the potential use of *WAPO1* in hexaploid wheat for further yield increases. The impact of *WAPO1* and SPS on yield depends on other genetic and environmental factors, hence, will require a finely balanced expression level to avoid the development of detrimental pleiotropic phenotypes.

## Introduction

Grain yield is a highly complex trait which is the product of a network of inter-connected yield-component traits, such as tiller number, inflorescence architecture and the rate of grain filling, often with interdependencies and trade-offs involved^1^. Studying individual yield-component traits, particularly those with higher heritability such as spikelet number per spike (SPS)^2^ can assist in reducing some of this complexity. As spikelets are specialized branches that produce grains, breeding for increased SPS could increase grain yield by increasing grain number and sink capacity.

SPS is affected by genes affecting flowering time through vernalization, photoperiod or earliness *per se (eps)* pathways: *VRN1, FUL2, FUL3, PPD1, VRN3*/*FT1, FT2, ELF3*^3–7^. Spring or winter growth habit is determined primarily by variation in the *VRN* genes. *PPD1* controls photoperiod-dependent floral induction, with the photoperiod insensitive allele *PPD-D1a* accelerating flowering, whilst *ELF3* is a minor effect *eps* gene. However, identification and characterization of major effect genes that increase SPS independently of these pathways has been limited so far.

The genetic dissection of grain yield and yield-component traits in hexaploid wheat is complicated by the compensatory effects of homoeologous genes. Their analysis in diploid model plants, such as rice, usually reveal QTLs with much higher effect sizes, e.g. 20-40% in rice vs 2-9% in hexaploid wheat^8^. Previous rice studies identified several genes involved in the regulation of panicle size and grain number and thus their orthologs provide a basis for studies in other cereals^9^. The rice gene *ABERRANT PANICLE ORGANIZATION 1* (*OsAPO1)* regulates panicle size, grain number^10,11^ and yield^12,13^ and higher expression of the *OsAPO1* transcript leads to more primary branches^11^. *OsAPO1* is syntenic with a major wheat SPS Quantitative Trait Locus (QTL) on chromosome 7AL in several bi-parental mapping populations and diversity panels^2,14–21^ as well as a minor homoeologous QTL on 7BL^22,23^.

More recently the *WHEAT ORTHOLOGUE OF APO1 (WAPO1)* has been proposed as the candidate gene for the SPS QTL on 7AL by multiple authors^24–26^, and validated in tetraploid wheat using a combination of mutants and transgenic lines^27^. However, transgenic overexpression of *WAPO-A1* in multiple backgrounds of hexaploid wheat has not yet been reported, limiting our understanding of gene function, effect size, and pleiotropic effects for the more relevant hexaploid wheat breeding.

Multi-parental Advanced Genetic Inter-Crossed (MAGIC) populations are well suited for QTL mapping due to their high levels of allelic diversity and recombination^28–30^. Based on residual heterozygosity in the MAGIC populations, QTLs can be further validated and fine mapped through the development of Heterogeneous Inbred Families (HIFs). Selected HIFs will be isogenic at most loci in the genome, but still heterozygous for the region spanning the QTL of interest. The resulting segregating progeny will have properties similar to near-isogenic lines (hereafter named NILs) with and without the QTL of interest and are ideal genetic material to perform precise phenotyping and transcriptomic analysis, and to assess the allelic contributions to the trait of interest as well as for fine mapping and candidate causal gene identification^31^.

In this study, we describe the functional validation of *WAPO1* in determining SPS in hexaploid wheat. We used two MAGIC populations across 14 site^×^year environments to identify two homoeologous SPS QTLs for 7AL and 7BL. Sets of contrasting NILs derived from HIFs segregating for the QTL on 7AL were used to validate and fine map the 7AL QTL, delivering a candidate gene *WAPO-A1* and its homoeologue *WAPO-B1*. For *WAPO-A1* we observed that increased expression in the spike in the NIAB MAGIC founder parents was positively correlated with SPS in the field. Therefore, our functional validation of the *WAPO-A1* gene in hexaploid wheat used a transgenic approach in both low- and high-SPS genetic backgrounds. This transgenic overexpression approach provided functional validation in hexaploid wheat for the role of *WAPO-A1* expression level in determining the number of SPS. Furthermore, we demonstrate that the increase in SPS in the transgenic lines is not always translated into an increased number of seeds per spike and can lead to pleiotropic or compensation effects under our growing conditions. The latter highlights the need to finely balance expression levels to achieve positive effects on yield when applied in breeding.

## Results

### The CSIRO and NIAB MAGIC populations have a wide phenotypic spread in SPS

Twelve field trials using an 4-parent MAGIC population^32^ from CSIRO were performed between 2011 and 2014 across three field sites in New South Wales, Australia (Yanco, Narrabri, Wallendbeen) and in a Greenhouse in Canberra (ACT, Australia). Two field trials using an 8-parent MAGIC population^28^ from NIAB were performed in 2014 and 2015 in Cambridge, UK. Heritability of SPS across all trials and both populations was high, varying from 0.45 to 0.86 with a mean of 0.71 (Supplementary Table 1).

Both MAGIC populations showed transgressive segregation for SPS. In the 4-parent CSIRO population, recombinant inbred lines (RILs) had a mean SPS number ranging from 12.6 to 25.0 compared to 16.3 to 23.3 for the MAGIC parents. Yitpi and Chara were identified as having a reduced mean number of SPS when compared to Westonia and Baxter. On average the difference was at least one spikelet per spike between the CSIRO parents in the low and high SPS groups. The 8-parent NIAB population, RILs had mean SPS number ranging from 17.70 to 29.80 compared to 20.32 to 26.40 for the MAGIC parents. The NIAB MAGIC parents could also be broadly divided into high and low phenotype groups, with Soissons, Robigus and Brompton having a reduced SPS number compared to the other five MAGIC parents (Alchemy, Claire, Hereward, Rialto, Xi-19).

### QTL mapping identified ten major SPS QTLs

QTL mapping was conducted using MPWGAIM^33^ with a separate univariate analysis for each trial. In total, 203 marker-trait associations (MTAs) for SPS were identified across 12 trials for the CSIRO MAGIC population and 18 MTAs for SPS were identified across 2 trials for the NIAB MAGIC population (QTL significance p < 0.05) (Supplementary Tables 2&3). The CSIRO MTAs cluster into ten major QTLs that are highly significant (max. -log10(p) > 5) and reproducible across at least two trials. For six of the identified QTL, we were able to assign candidate genes based on previously known allelic effects of flowering pathway genes, leaving four QTL that were initially not annotated. Of these, a SPS QTL on 7AL was observed most consistently at a high significance and was chosen for further downstream analysis (Table 1).

**Table 1:**
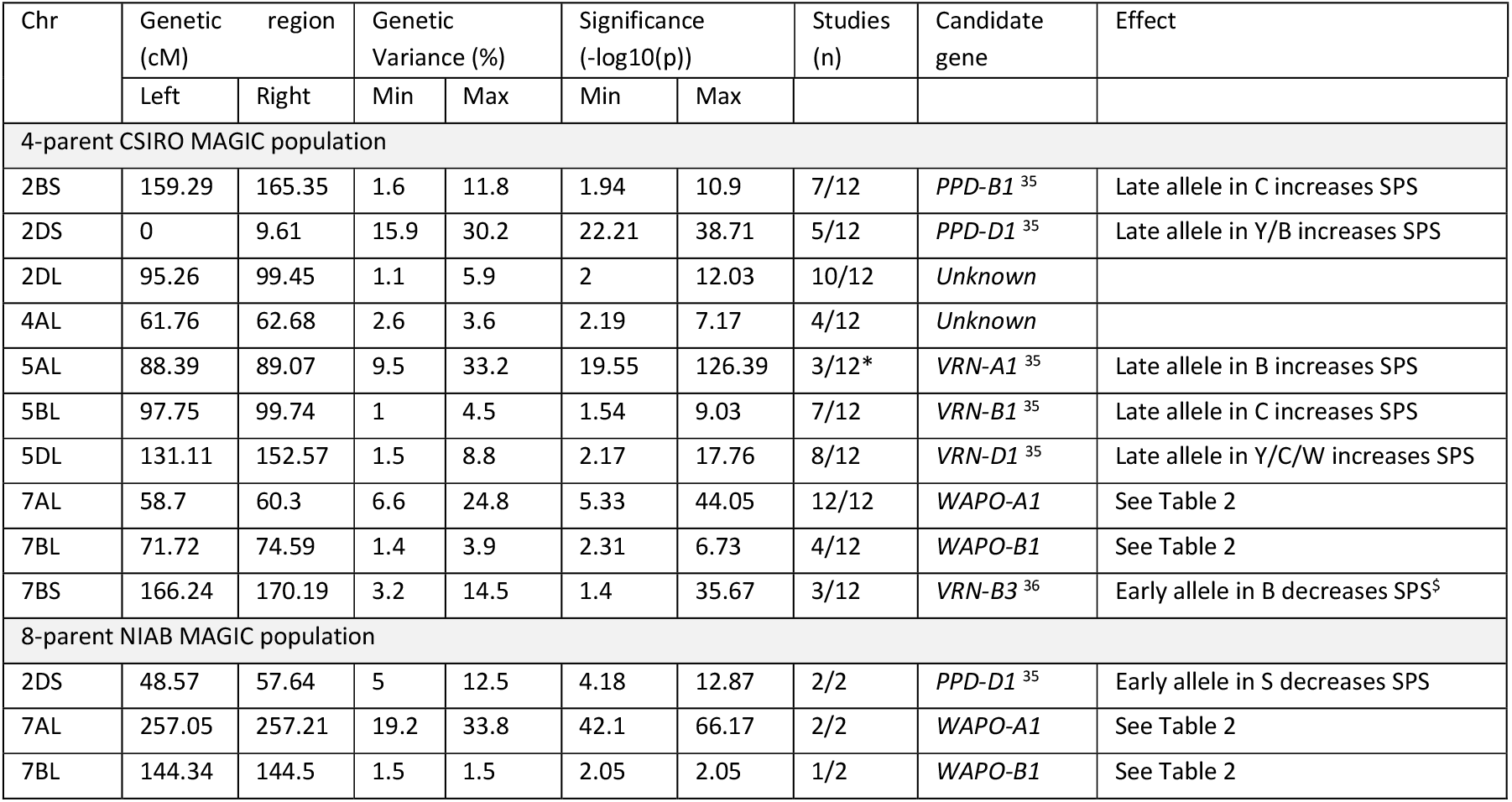
Overview of major SPS QTLs in the 4-parent CSIRO and the 8-parent NIAB MAGIC populations and the underlying candidate genes. QTLs for SPS from 12 trials with the 4-parent CSIRO MAGIC population and two trials with the 8-parent NIAB population. QTL analyses were conducted using whole genome average interval mapping using MPWGAIM^33^. For the NIAB MAGIC population a genetic map adapted from Gardner et al., 2016^34^ was used. For the CSIRO MAGIC population a previously published map was used^33^. Genetic variance (%) and significance (-log10(p)) as calculated by MPWGAIM^33^. *The QTL likely associated with *VRN-A1* was only observed in the CSIRO Greenhouse studies, where vernalization did not occur. ^$^Inferred from reported *VRN-B3* allele. Variety abbreviations: Baxter (B), Chara (C), S (Soissons), W (Westonia), Y (Yitpi)

Despite substantially different environmental conditions, the major QTL for SPS on 7AL was observed in all 12 CSIRO trials and in two NIAB trials. This indicates that this QTL was environmentally and genetically very stable (Supplementary Table 4). In the 4-parent CSIRO population, the 7AL SPS QTL defined a 5.7Mb interval between 669.7Mb and 675.4Mb on the IWGSCv1.1 assembly, with a global significance level of -log10(p)>5 across all 12 trials and mean genotypic variance explained of 13.6%. Baxter and Westonia carry the high SPS genotype, with an average increase of 1.1 spikelets per spike in the MAGIC RILs compared to the low SPS genotype (Yitpi, Chara), under these conditions. In the 8-parent NIAB population the 7AL SPS QTL defined a narrower 2.3Mb interval between 672.0Mb and 674.3Mb. In the 2014 trial, the QTL explains a remarkable 33.8% of genetic variation in SPS within the NIAB 8-way MAGIC population at a global significance level of -log10(p) 66.17. Alchemy, Claire, Hereward, Rialto, Soissons, Xi-19 all carry the high SPS genotype, with an average increase of 2.1 spikelets per spike in the MAGIC RILs compared to the low SPS genotypes (Brompton, Robigus).

In addition to the major 7AL QTL for SPS, a homoeologous SPS QTL on 7BL with a smaller effect size was also identified in both the 4-parent CSIRO and 8-parent NIAB populations (Table 1). In the 8-parent NIAB population, this second QTL is located on a 4.9Mb interval (645.3 – 650.1Mb) directly homoeologous to the 7AL QTL (-log10(p) = 2, Genetic Variance Explained (GVE) = 1.5%). Claire, Xi-19 and Soissons carry the high 7BL SPS parental genotype. Four of the 4-parent CSIRO trials also had a 7BL QTL mapping in the vicinity (642.5Mb – 655.0Mb) with Yitpi and Baxter carrying a high SPS parental genotype (-log10(p)>2.3, mean GVE = 1.5%).

### Validation and fine mapping of the 7A SPS QTL using HIFs

To validate and fine-map the 7AL QTL, four Heterogenous Inbred Families (HIFs) were derived from selected F_7_ RIL lines of the 4-parent CSIRO population containing heterozygosity in the QTL region. These RILs were utilized to develop sets of NILs with contrasting presence of the 7AL SPS QTL.

Four NIL pairs (F0035, F0748, F1229, and F1570) showed consistent differences in SPS between the contrasting alleles across the 7AL locus, thus validating the QTL (Figure 1A).

**Figure 1:**
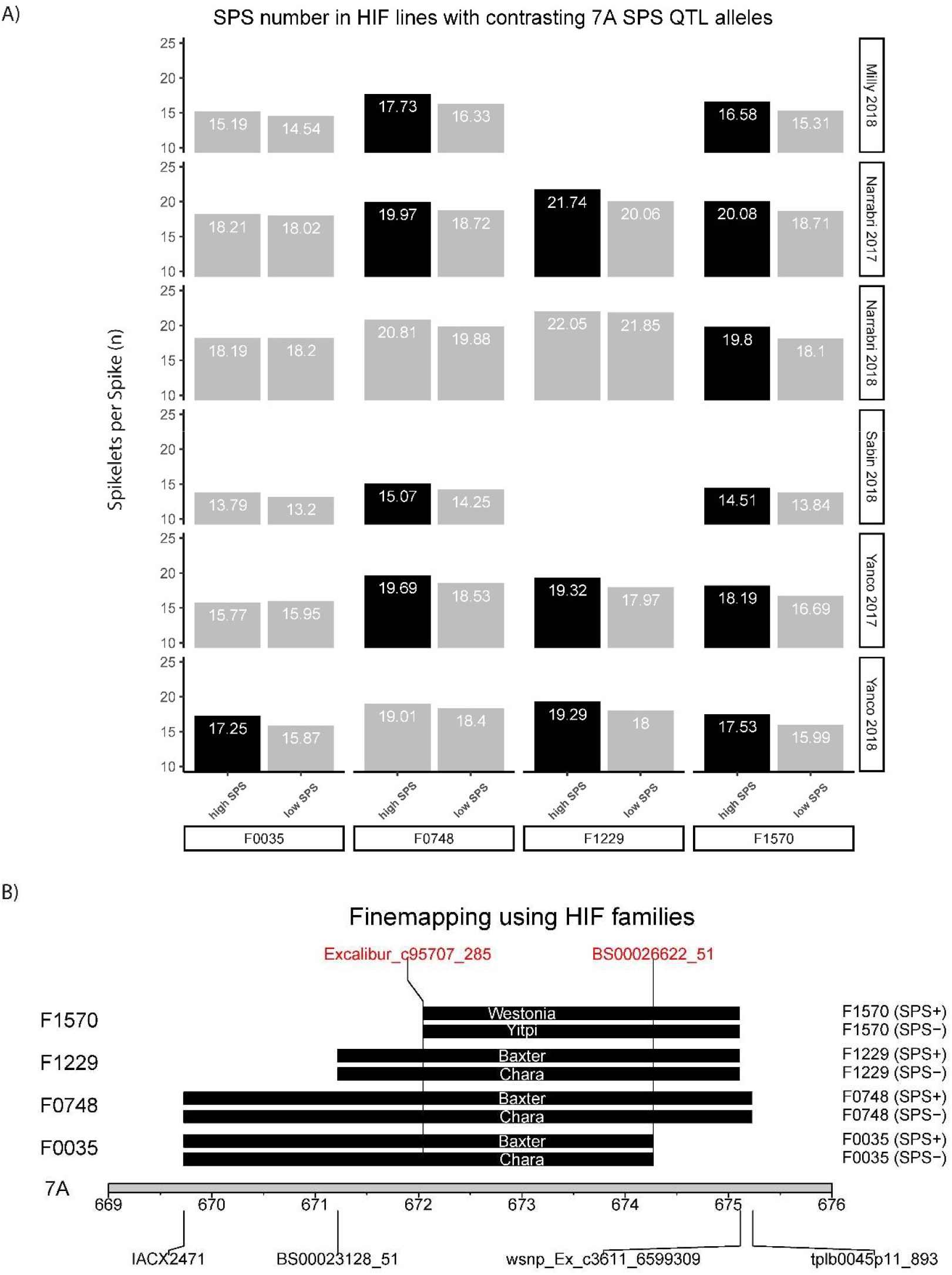
Validation and fine mapping of the 7A SPS QTL using HIF families derived from the CSIRO 4-way MAGIC population. A) Summary of differences in mean SPS values across NIL pairs for the 7AL SPS QTL. Black fill indicates a significant increase (p<0.05) between NIL pairs. B) Finemapping using HIF families. Contrasting founder genotypes in the NIL pairs are shown. The common QTL region is delimited by markers Excalibur_c95707_285 and BS00026622_51. The NIL pairs were phenotyped at four locations in France, the USA and Australia during 2017 and 2018 (6 site^x^year combinations).

For two NIL pairs (F0748 and F1570) nearly all differences were statistically significant (p < 0.05). Because the selected NILs carried the 7AL QTL genotypes of all four founder parents, the allelic effects for each of the founders of the 4-parent CSIRO population could be determined. In addition, the SPS region was further delimited from 5.7Mb to an interval of 2.2Mb by NIL pairs F0035 and F1570 (Figure 1B).

Additional phenotyping revealed the high SPS lines in each NIL pair also showed an increase in Grain number per spike in most cases, albeit mostly insignificant, (Supplementary Figure S1) and no differences in heading date exceeding 0.5 days were observed (Supplementary Figure S2).

### Identification of *WAPO-A1* and *WAPO-B1* as homoeologous candidate genes for SPS

The QTL region delineated using the HIFs on 7AL contained twenty genes. Both the 7AL and 7BL QTL identified in the NIAB and CSIRO MAGIC populations are syntenic to rice chromosome 6 (Figure 2). Within these three homoeologous or syntenic QTL regions, four conserved orthologous protein groups were identified: *TraesCS7A02G480100/TraesCS7B02G382300/OsHMA9, TraesCS7A02G481500/TraesCS7B02G383900/OsAAH, TraesCS7A02G481600/TraesCS7B02G384000/OsAPO1* and *TraesCS7A02G482000/TraesCS7B02G384200/Os06g0665100*.

**Figure 2:**
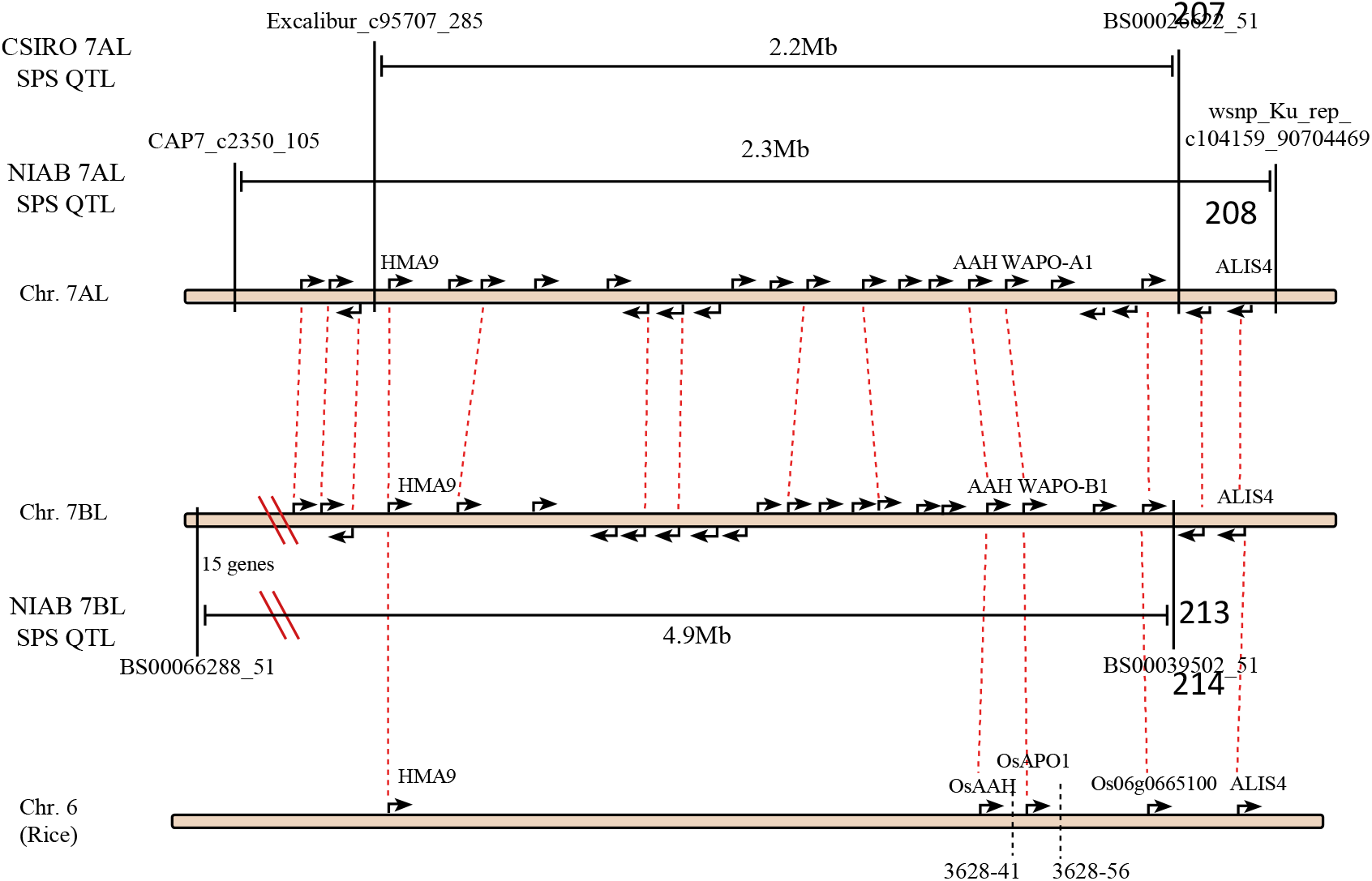
Syntenic relationships between the 7AL and 7BL SPS QTLs in the 4-parent CSIRO MAGIC population and the 8-parent NIAB MAGIC population and the rice SCM2/qPBN6 QTL^12,39^. Arrows indicate locations of IWGSCv1.1. gene models obtained from Ensembl Plants release 51. Dashed red lines indicate homoeologous and orthologous relationships as defined by Ensembl Plants 51. Gene descriptions from Ensembl Plants 51 have been added where available. Physical distances are not drawn to scale.

*OsHMA9* is a metal-efflux protein and *OsAAH* an allantoate deiminase. Neither have previously been linked to SPS phenotypes^37,38^.

*OsAPO1* is an F-box protein that had previously been shown to improve rice yield, primary rachis branching and lodging resistance as the causal gene of *qPBN6*^39,40^ and *SCM2* QTL in rice^12^. The wheat orthologues of *OsAPO1* on 7AL and 7BL (*WAPO-A1* and *WAPO-B1*) were thus identified as good candidate genes for the 7AL and 7BL SPS QTLs. We also confirmed that no SPS QTLs were detected in the *WAPO-D1* genomic region. *WAPO-A1* and *WAPO-B1* genes and *WAPO-A1* promoter sequences for Fielder, Chinese Spring, Robigus and Claire were obtained from public resources^41–43^. *WAPO-A1* and 5kb of upstream promoter sequence in Yitpi, Baxter, Chara and Westonia were obtained by Sanger Sequencing (GenBank ON210994-ON210998).

Sequence alignments showed that the high SPS *WAPO-A1* allele carries a 115bp promoter deletion and two amino acid changes (C47F and D384N). Based on recent *WAPO-A1* nomenclature^24^ we reclassified our alleles accordingly (Table 2). All the assigned *WAPO-A1* alleles are consistent with the observed founder effects in both MAGIC populations for the 7AL SPS QTL.

We also classified *WAPO-B1* alleles based on recent nomenclature^22^ (Table 2). All assigned *WAPO-B1* alleles are also consistent with the observed founder effects except for Soissons, which has a high SPS founder effect on 7BL, while carrying the low SPS *WAPO-B1*.*hap1* allele. Westonia carries the *WAPO-B1*.*hap3* allele, previously reported to carry a frameshift mutation/premature stop codon^22^. Based on our alignments and the protein sequence of previous *WAPO-B1* gene model TRIAE_CS42_7BL_TGACv1_578478_AA1895640.1 we believe that this gene model is annotated incorrectly (Supplementary Figure S3) and that as a result, this InDel is unlikely to cause a premature stop as it falls into the 5’ UTR of *WAPO-B1*. This observation would be consistent with the fact that *WAPO-B1*.*hap3* only has an intermediate SPS phenotype across four trials.

**Table 2:** Founder effects and *WAPO-A1* and *WAPO-B1* alleles of NIAB and CSIRO MAGIC parents. Founder effects classified based on MPWGAIM predictions. *WAPO-A1* and *WAPO-B1* alleles classified according to Kuzay et al., 2019^24^ and Corsi et al., 2021^22^. *Founder phenotypic effects for Fielder and Chinese Spring (CS) are inferred based on genome sequence information since they are not MAGIC founder parents. Detailed variant calls presented in Supplementary Table S5.

| Variety | 7A SPS QTL |  | 7B SPS QTL |  |
| --- | --- | --- | --- | --- |
|  | Founder effect at 7AL | *WAPO-A1* allele | Founder effect at 7BL | *WAPO-B1* allele |
| Yitpi | low SPS | *WAPO-A1a* | high SPS | *WAPO-B1.h2* |
| Baxter | high SPS | *WAPO-A1b* | high SPS | *WAPO-B1.h2* |
| Chara | low SPS | *WAPO-A1a* | low SPS | *WAPO-B1.h1* |
| Westonia | high SPS | *WAPO-A1b* | intermediate SPS | *WAPO-B1.h3* |
| Alchemy | high SPS | *WAPO-A1b* | low SPS | *WAPO-B1.h1* |
| Brompton | low SPS | *WAPO-A1a* | low SPS | *WAPO-B1.h1* |
| Claire | high SPS | *WAPO-A1b* | high SPS | *WAPO-B1.h2* |
| Hereward | high SPS | *WAPO-A1b* | low SPS | *WAPO-B1.h1* |
| Rialto | high SPS | *WAPO-A1b* | low SPS | *WAPO-B1.h1* |
| Robigus | low SPS | *WAPO-A1a* | low SPS | *WAPO-B1.h1* |
| Soissons | high SPS | *WAPO-A1b* | high SPS | *WAPO-B1.h1* |
| Xi-19 | high SPS | *WAPO-A1b* | high SPS | *WAPO-B1.h2* |
| CS* | high SPS | *WAPO-A1b* | high SPS | *WAPO-B1.h2* |
| Fielder* | high SPS | *WAPO-A1b* | low SPS | *WAPO-B1.h1* |

### *WAPO-A1* expression in the field is correlated with SPS

To investigate if the expression of *WAPO-A1* in wheat is correlated with SPS in the field we collected samples from tiller dissections of the eight NIAB MAGIC parents from a 2017 field nursery at growth stage GS32^44^, because wheat-expression.com^45^ data suggests that *WAPO1* homoeologous are expressed in the spike at this developmental stage. Expression analysis using qRT-PCR confirmed that higher *WAPO-A1* expression in wheat inflorescences is correlated (R^2^ = 0.77) with increased SPS (Figure 3). The low expression of *WAPO-A1* in Soissons may be a result of the presence of the *Ppd-D1a* allele in this variety, which accelerates flowering time, so we possibly missed the expression peak of *WAPO-A1*.

**Figure 3:**
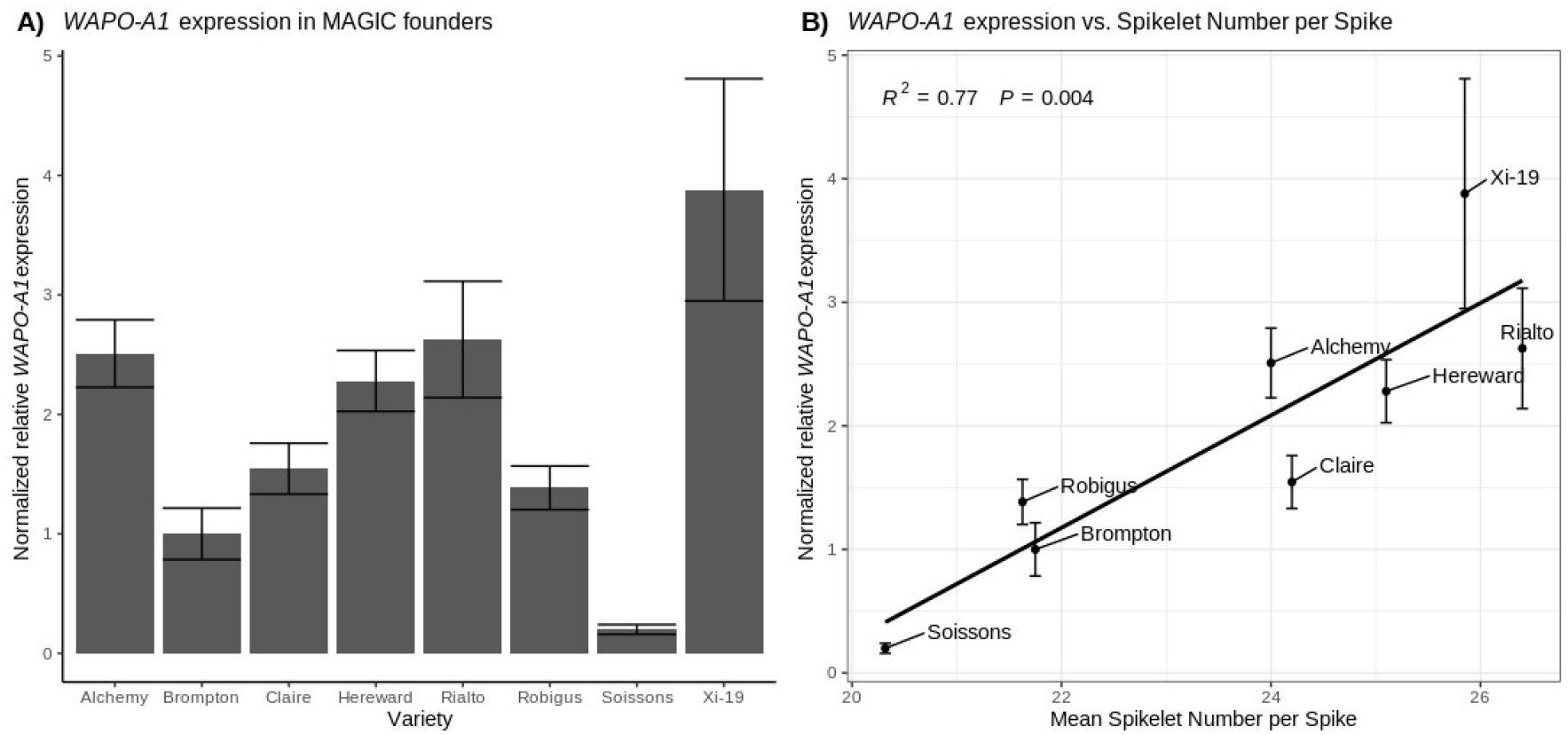
*WAPO-A1* expression is correlated with spikelet number per spike. All samples were collected at GS32 apart from Soissons which at the collection date had advanced to GS34. A) Expression of *WAPO-A1* transcript relative to the housekeeping genes *TaRP15*^46^ and *Ta2291*^47^and normalized to *WAPO-A1* expression in Brompton. B) Regression of expression of *WAPO-A1* (n = 3) on mean SPS for the MAGIC Founder lines (n=20) in the 2014 field trial. Error bars are SEM.

### Transgenic complementation of the *WAPO-A1a* allele with *WAPO-A1b* leads to increased SPS and spike length

To functionally validate the *WAPO-A1* candidate, Yitpi, carrying the low SPS *WAPO-A1a* allele, was complemented with the native high SPS *WAPO-A1b* allele and promoter from Westonia (∼8Kb genomic fragment, see methods) using Agrobacterium mediated transformation. Six sets of T_2_ single-insertion homozygous lines next to their respective null-segregant control line were grown in the glasshouse and phenotyped (Figure 4A, 4B and Supplementary Figure S4).

**Figure 4:**
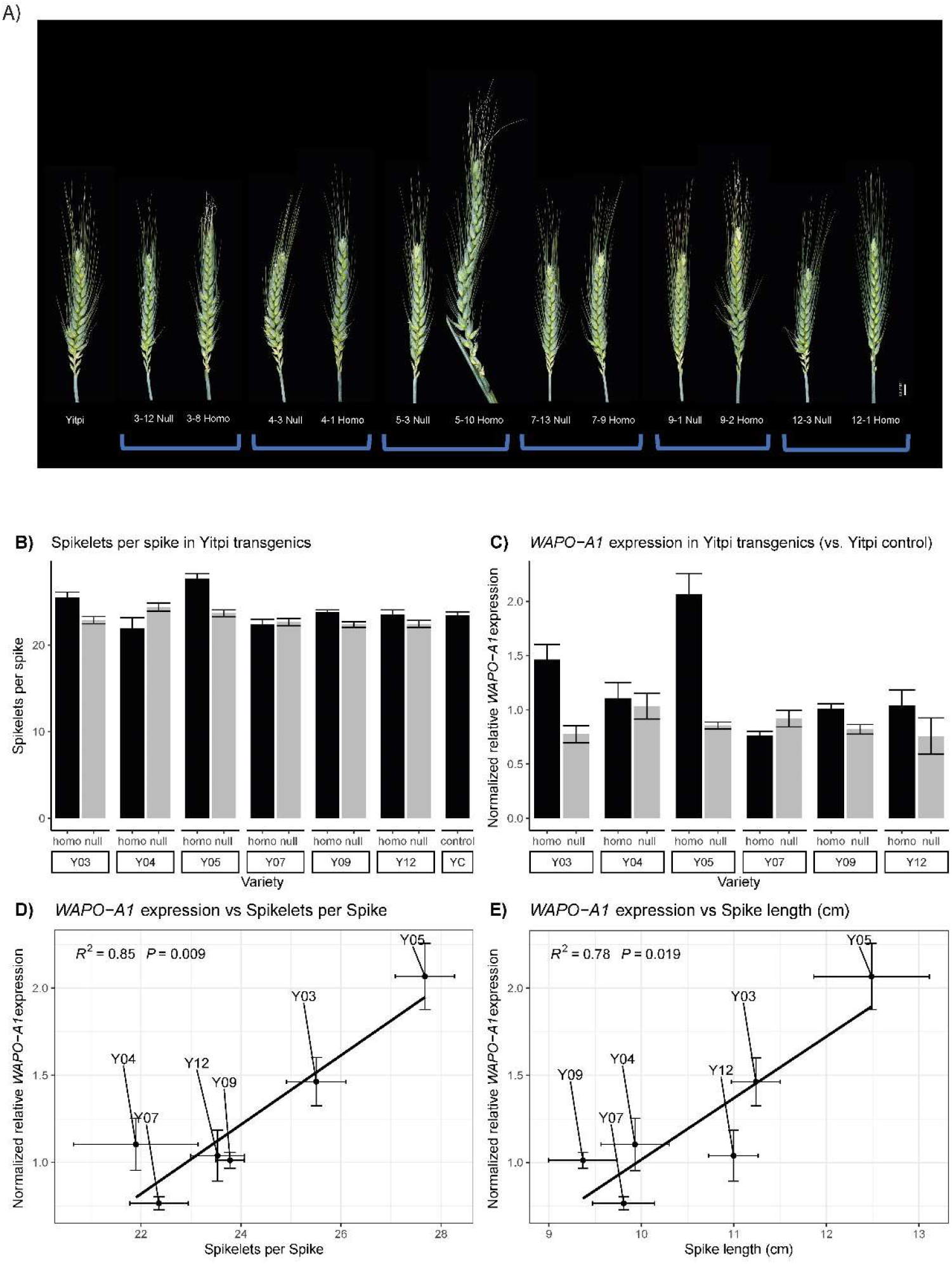
Spike phenotypes and WAPO-A1 expression in Yitpi transgenic and control lines. A) Representative spikes from transgenic lines, their null-segregants and non-transformed Yitpi plants. We observed the appearance of infertile basal spikelets in all genotypes although they were more common in transgenic plants. The presence of paired-spikelets was similar across all type of plants. B) Mean SPS number phenotyped at maturity in transgenic lines, their null-segregants and non-transformed Yitpi plants. C) Gene expression was assayed in developing spikes dissected at GS32 and normalized to the Yitpi WT control. The means and SEM of three biological replicates are shown. D) Regression of expression of *WAPO-A1* on SPS for the six transgenic lines carrying the Westonia *WAPO-A1* transgene. E) Regression of *WAPO-A1* expression on Spike length. For SPS and Spike length, 50 spikes (5 spikes x 10 plants) of each genotype were analysed

Four out of six transgenic lines showed an increase in SPS compared to their null segregant. Homozygous T_2_ lines #5 and #3 displayed the strongest phenotype reaching more than 27 and 25 SPS, respectively, as compared to 23 in their null segregants. Homozygous plants from lines #9 and #12 displayed a gain of about one SPS, while lines #4 and #7 did not show a significant difference in SPS compared to their null segregants. The number of SPS in a non-transformed Yitpi control was similar to the null-segregant lines (Figure 4B).

Samples of developing spikes at GS32 were collected from these lines for gene expression analysis using qRT-PCR (Figure 4C), which revealed a strong correlation between the level of *WAPO-A1* expression and the number of SPS and spike length (Figure 4D, E). These results are consistent with a causal, positive correlation between *WAPO-A1* expression and the number of SPS and spike length.

### SPS levels can be increased above those conferred by the native *WAPO-A1b* allele

To determine if SPS and seeds per spikelet could be increased beyond the levels of the high SPS *WAPO-A1b* allele, we also transformed Fielder, which already contains a native high-SPS *WAPO-A1b* allele, with the native high-SPS *WAPO-A1b* allele and promoter from Westonia. We generated five independent primary transgenics in the Fielder background and isolated three single-insertion lines, for which we identified homozygous T_2_ and null-segregants for phenotypic analysis of SPS, empty basal spikelets and seeds per spike.

All three transgenic lines showed an increased number of SPS in comparison to the nulls and the number of SPS was strongly correlated with the level of transgene expression (Figure 5, Supplementary Figure S5). These results further validated the conclusion that SPS number is closely correlated with WAPO-A1 expression levels. It also demonstrates that it is possible to increase SPS number beyond the levels determined by the native high SPS allele alone.

**Figure 5:**
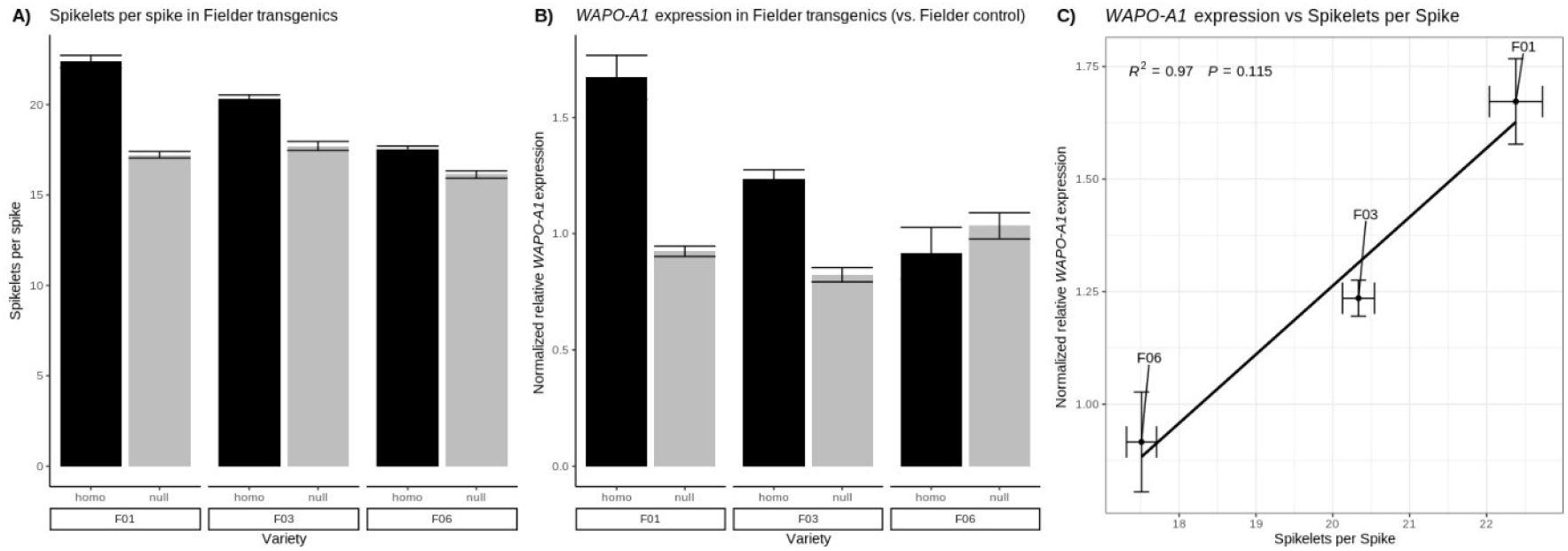
WAPO-A1 expression analysis in Fielder transgenic and control lines. A) Mean SPS of transgenic and null-segregants of plants phenotyped at maturity; B) Gene expression was assayed in developing spikes dissected at GS32. Expression is normalized to the Fielder WT control. C) Regression of expression of *WAPO-A1* on SPS for the transgenic lines. Means and SEM of three biological replicates are shown.

The Fielder control lines have fewer SPS (SPS = 17.02) than the Yitpi control (SPS = 23.44), despite carrying the favourable *WAPO-A1b* allele. This difference in SPS between the two cultivars could be explained by different photoperiod responses. Fielder carries the photoperiod insensitive *PPD-D1a* allele while Yitpi carries the photoperiod sensitive *PPD-D1d* allele, known to increase SPS^4^.

Importantly, all the homozygous transgenic lines of Fielder, but not Yitpi containing the native high SPS *WAPO-A1b* transgene had spikes displaying a “compact head” phenotype (Supplementary Figure S6), which was also previously described^27^. This phenotype was observed in all three Fielder families in the homozygote T_2_ lines but was not seen in the null-segregants.

### Increases in *WAPO-A1* expression may cause pleiotropic effects

In hexaploid wheat an increase in infertile spikelets in *WAPO-A1b* lines compared to *WAPO-A1a* has previously been described^22,26^. In Kronos, a tetraploid wheat, a reduction in spike fertility has also been reported^27^, in lines containing additional WAPO-A1 transgene copies. In our studies the transgenic lines with higher levels of *WAPO-A1* expression show an increased number of empty basal spikelets (Supplementary Figure S7). This was especially pronounced in the highest expressing lines, both in Yitpi (T_2_ line #5) and Fielder (T_2_ line #1). Similar to phenotypes previously described in tetraploid wheat (Kronos) *WAPO-A1* overexpression lines^27^ we observed one or two malformed florets at the base of some spikes of the transgenic lines. The phenotypic alterations are indicative for developmental disorders like homeotic transformations or failed organ development (Supplementary Figure S8).

In rice, lines with increased *OsAPO1* expression reveal a suppression of tiller outgrowth^11^. Similarly, we observed a negative association between tiller number and *WAPO-A1* expression across our transgenic Yitpi lines in the greenhouse pot experiment (Supplementary Figure S7). However, the null segregants also showed variability in this trait meaning the decrease was only significant in one line (T_2_ line #5).

### Increased SPS in transgenic lines does not translate into equivalent increases in seed yield

Under our greenhouse conditions we could not measure a clear benefit in yield (as seeds per spike or per plant) associated with transgene expression (Supplementary Figure S4), either due to the pleiotropic phenotypes described above or potential source limitations under these growth conditions.

The transgenic Yitpi line with the highest transgene expression and number of SPS (T_2_ line #5, T_2_ line #3) showed a clear penalty in both seeds per spike and seeds per plant. We also observed an extreme branching phenotype and floral defects in a small proportion of the spikes developed by Yitpi T_2_ line #5 (Supplementary Figure S8). In fact, the spikes were often so long that they did not fully emerge from the underlying leaf sheath, and the florets inside were often infertile (Figure 4A). Taken together, our data indicate, that a too high expression of *WAPO-A1* reveals a rather negative effect on seed production.

However, lines with a lower increase of the transgene and a resulting smaller increase in SPS such as Yitpi T_2_ Line #12 (*WAPO-A1* expression increase 37%; Seeds per spike increase 11%) and Fielder T_2_ line #3 (*WAPO-A1* expression increase 50%; Seeds per spike increase 21%) indicated that increases in seeds per spike could be observed, suggesting that one can select an optimal range of *WAPO-A1* expression levels to achieve the desired phenotypic and yield effects.

## Discussion

### MAGIC populations are well suited for QTL dissection

The use of two separate MAGIC populations across 14 site^×^year environments enabled us to capture a wide range of genetic and phenotypic variation and to identify homoeologous SPS QTL on 7AL and 7BL. The additional recombination present in MAGIC populations allowed for improved mapping resolution which aided the direct identification of candidate genes, as was the case with the NIAB MAGIC population, or via the generation of HIF lines^31^. This pipeline for candidate gene discovery via HIF lines derived from MAGIC populations is repeatable using other yield-component traits.

MAGIC populations can also reveal the effects of stacking of favourable alleles through transgressive segregation^48^ as well as epistatic effects^49^. Within the 4-parent CSIRO population alleles at the *Ppd-D1, PPD-B1, VRN-A1, VRN-B1* and *VRN-D1* loci are segregating in addition to *WAPO-A1* and *WAPO-B1*^32,35^. In addition to *WAPO-A1* and *WAPO-B1*, within the 8-parent NIAB population *Ppd-D1, ELF3-B1* and *ELF3-D1* are also segregating ^4,7,28,50^. All of these loci were identified as significant SPS QTL in the respective populations with the exception of *ELF3-B1* and *ELF3-D1*, which may be due to their minor effect on SPS or due to more complex environmental interactions of *ELF3*^51^.

### Overexpression of *WAPO-A1* in hexaploid wheat increases SPS

The *WAPO-A1* gene has been proposed as the candidate gene for a major QTL for SPS on 7AL mapped in several populations^22–26^ and the *WAPO-B1* gene as a candidate for a smaller QTL for SPS on 7BL^22^. Recently, ^27^Kuzay et al., 2022 demonstrated, using CRISPR-generated mutants and transgenic plants, the role of *WAPO-A1* and *WAPO-B1* in determining SPS in tetraploid wheat.

Here, we provide functional evidence in hexaploid wheat for a similar role of the *WAPO-A1* gene in determining SPS in a dose dependent manner. We have generated two sets of transgenic lines to test the effect on supplementing cultivars carrying either the low-SPS *WAPO-A1a* or the high-SPS *WAPO-A1b* alleles with a *WAPO-A1b* allele from a high-SPS cultivar (Westonia). In both sets of transgenic lines, we observed a positive correlation between *WAPO-A1* expression and the number of SPS, which is consistent with variation in *WAPO-A1* being responsible for the observed QTL. ^27^Kuzay et al., 2022 provided strong evidence for a role of *WAPO-B1* in determining SPS, by a reduced number of SPS present in *wapo-b1* mutants. However, it was previously reported that overexpression of *WAPO-D1* in the elite hexaploid variety KN199 had no effect on spike phenotypes^19^. Different *WAPO1* homoeologues may therefore have different effects on SPS.

At the molecular level, *WAPO-A1* increases the expression of B-Class and C-Class MADS box genes, which affect lodicules, stamen and carpel development^52^. This is a possible cause behind the floral abnormalities and reduced fertility observed in both the transgenic Yitpi and Fielder lines. In the Fielder transgenics we also observed a compact spike phenotype, which indicates that the overexpression of *WAPO-A1* in these lines alters spike development in several ways. Similar phenotypes in the florets and spikes have been reported in transgenic lines overexpressing the *WAPO-A1* gene in the tetraploid wheat cv Kronos^27^.

In transgenic lines displaying a low level of expression of the *WAPO-A1* transgene, and a small increase in SPS, we observed an increase in the number of seeds per spike (Yitpi line #12 and Fielder line #3). This is in line with the idea that a subtle rather than a strong increase of *WAPO-A1* expression levels could be associated with increased grain number. This is similar to the situation in rice in which *SCM2*, a weaker allele of *OsAPO1* compared to *Ur1*, has been reported to increase spikelet numbers without reducing panicle number^12^. For commercial application of transgenic lines, further validation of the positive yield component phenotypes observed in transgenic lines with modest increases in *WAPO-A1* expression in a replicated field trial environment will be required. The long timelines, high costs and current lack of commercial transgenic traits in wheat also represent significant challenges for product development.

### *WAPO-A1* activity is finetuned via interconnected, developmental pathways

QTL mapping in the two MAGIC populations revealed large phenotypic effects of *PPD-D1a* and *VRN-A1a* (and likely *VRN-B3* as well) on the number of SPS. More minor but significant effects on SPS from *PPD-B1a, VRN-B1a* and *VRN-D1a* were also observed. The MAGIC populations studied here thus form an excellent platform to identify the genetic backgrounds that optimise the SPS, and grain number potential associated with *WAPO-A1b* for specific environments.

*PPD1, VRN1* and *VRN3* all influence the timing of transition from vegetative to inflorescence meristem (IM) and from IM to terminal spikelet^3–5,53^. Later transitions are associated with increased SPS^53^, either linked to the duration or resources available for development. This would explain the low SPS observed for Soissons, carrying the early *PPD-D1a* allele, despite having the high-SPS *WAPO-A1b* allele. Similarly, in our greenhouse experiments Fielder (*WAPO-A1b, PPD-D1a*) also had fewer SPS than Yitpi (*WAPO-A1a, PPD-D1d*). *WAPO-A1* native alleles on the other hand have not been reported to cause large changes in flowering time in field trials^25,53^ and while transgenic expression delayed flowering, CRISPR and EMS mutants had no significant effects^27^.

Our results show that the *WAPO-A1b* allele has a larger effect on the number of SPS in the winter wheat NIAB MAGIC population (+2.1 SPS) compared to the spring wheat CSIRO MAGIC population (+1.1 SPS; *VRN-A1a* and/or *VRN-B1a* and/or *VRN-D1a* present, see Table 1). An epistatic interaction between *WAPO-A1, VRN-D3, PPD-B1* and *Qsn*.*csu-6B* has recently been described^53^. These data together support the conclusion that the phenotypic effect of *WAPO-A1* alleles on SPS is finetuned by the timing of developmental transitions and the genetic constitution of *PPD1, VRN1 and VRN3*.

The interactions of *WAPO-A1b* with flowering pathways can also be discussed at the molecular level. *PPD1, VRN1* and *VRN3*/*FT* form part of a signalling cascade that leads to the upregulation of *SOC1-1* and *LFY*^54^. The orthologue of *LFY* is *OsAPO2*^55^ in rice and *HvLEAFY* in Barley^56^, which interact with the *WAPO1* orthologues *OsAPO1* or *HvUFO*, respectively. OsAPO1 and OsAPO2 form a regulatory module together with LARGE2^32^. *OsAPO2* has also been previously shown to be transcriptionally regulated by *SP3*^57^ in rice. Together the *WAPO1* pathway genes are expressed in the IM and lateral meristems and extend IM maturation and control spikelet meristem identity acquisition^58^. This may also explain the extreme branching phenotype observed in some spikes of the transgenic Yitpi T_2_ line #5.

To further elucidate the complexity of these regulatory networks, further work should focus on determining epistatic relationships between the identified QTLs at the phenotypic level as well as relating this to more comprehensive studies of expression changes and molecular interactions in different genetic backgrounds.

### *WAPO1* breeding applications

Increasing SPS is a potential path to increasing seeds per spike, grain number and ultimately yields. *PPD1* and *VRN1* have already been deployed to locally adapt cultivars to geographic adaptation zones and due to their importance for local adaptation it can be assumed that there would likely be extensive trade-offs in their use in improving SPS. *WAPO-A1b* on the other hand is potentially valuable for wheat breeders as it can increase SPS in locally adapted varieties without having strong impacts on flowering time.

However, the increased number of SPS found in most of the transgenic lines is not always translated in an increased number of seeds per spike. Perhaps, under our glasshouse growing conditions, the plants did not have sufficient resources to fill the extra spikelets, which might be especially relevant in pot experiments, where a large number of tillers and spikes per plant is formed as compared to the situation in dense canopies under field conditions. Additionally, strong overexpression of *WAPO-A1* produces some defects during floret or seed development.

These results align with those of other publications that report on the pleiotropic effects of *WAPO-A1b*, including reduced spike fertility^26^ and increased seeds per square meter but reduced TGW^15^. As previously proposed^24^, our results imply that *WAPO-A1b* promotes an increase in sink capacity which may require introgression into high-biomass productive varieties and cultivation under conditions that support high yield potential. The observed trade-offs are not unique to *WAPO1* and have been shown in other genes that increase sink capacity such as *FT2*^59^ and *FUL2*^3^. In rice it has also been reported that Os*APO1* increases yield^13^, but that the effect on yield is source dependent^60^ and dependent on gene expression levels as described above.

It might be possible to overcome these trade-offs by stacking *OsAPO1*/*WAPO-A1* with genes that have complementary phenotypic effects, and some promising examples have been reported in both rice and wheat. For example, stacking *WAPO-A1* with *GNI-A1* and *qGSNP-A-5A* to further increase SPS without reducing TGW was possible in a Japanese dihaploid wheat population^14^.

An alternative approach may be to exploit genes known to interact at the molecular level. In rice it was recently shown that that OsAPO1 physically interacts with OsAPO2 and a HECT-domain E3 ubiquitin ligase LARGE2/OsUPL2. LARGE2 modulates the protein stability of OsAPO1 and OsAPO2^61^. Extrapolating from these results in rice, further finetuning of WAPO1 activity and resulting SPS and seed number might be possible in wheat through posttranscriptional mechanisms and may help to overcome the constraints due to the limited number of *WAPO-A1* alleles that have currently been identified.

## Conclusion

This study has provided functional validation in hexaploid wheat for the role of *WAPO-A1* in determining SPS. We demonstrated that the expression level of this gene in MAGIC lines under field conditions as well as in transgenic lines in the greenhouse is positively correlated with the number of spikelets per spike. We also identified potential pleiotropic effects of excessive *WAPO-A1* expression. Future work can focus on identifying additional existing or novel *WAPO1* alleles that have an optimized level of expression and protein activity or identifying pathways and genes that can be used to optimize the expression of *WAPO1* or that have complementary phenotypic effects. The optimal activity of *WAPO-A1* resulting into yield gains can be expected to be dependent on the yield potential of the environment and agronomic practice.

## Methods

### Plant Materials

The founder parents for the NIAB and CSIRO wheat MAGIC populations were selected based on wide genotypic and phenotypic diversity.

The 4-parent CSIRO MAGIC population founders are spring wheat cultivars from Australia (Yitpi, Baxter, Chara and Westonia)^32^.

Six of the NIAB MAGIC population founder parents are winter wheat cultivars from the United Kingdom (UK) (Alchemy, Brompton, Claire, Hereward, Rialto, Robigus) and one from France (Soissons) and one is a facultative spring variety (Xi-19)^28^.

All plant material used in this study was derived from commercially released wheat varieties that were obtained and used in accordance with applicable institutional, national and/or international regulations.

### SPS Phenotyping

For the 4-parent CSIRO MAGIC population, F_6_ derived RILs were grown in partially replicated trials^62^ (partial replication rate of 1.2 - 1.5): 1080 lines were grown in a Greenhouse (Canberra, ACT, Australia, CSIRO Black Mountain Site; 2011(F_9_), 2012(F_10_), 2013(F_10_)), 1009, 960 and 1008 field plots were grown in Narrabri (New South Wales, Australia; 2012(F_10_), 2013(F_10 &_ F_11_), 2014(F_12_)), 1500, 1800 and 1009 field plots in Wallendbeen (New South Wales, Australia; 2011(F_9_), 2012(F_10_), 2013(F_10_)) and 1120, 1920 and 912 in Yanco (New South Wales, Australia; 2012(F_10_), 2013(F_10_), 2014(F_12_)). Three representative wheat ears were collected from each plot and the spikelets per spike (the total number of rachis nodes plus the terminal spikelet) were hand counted starting at the bottom of each spike.

Additionally, a fully replicated trial of 784 F_7_ 8-parent NIAB MAGIC RILs and the eight founders was grown at the NIAB experimental farm in Cambridge, UK during the 2013/2014 field season. Ten representative wheat ears were collected from 1000 of the 1600 plots in the field and dried at room temperature. Collection was done in a partially replicated design with 200 RILs and the MAGIC founders collected in duplicate. In 2014/2015 a nursery of 1091 F_8_ MAGIC lines and the founders was screened for SPS as described above by collection of six representative wheat ears.

Selected F_12_, F_13_ NIL sets were grown in 2017 and 2018 in Narrabri and Yanco, NSW, Australia and in 2018 in Sabin, Minnesota, USA and Milly-la-Forêt, France. In 2017 each line was grown in four replicates and in 2018 in six replicates. Six representative wheat ears were collected from each plot. Additional phenotyping for grains per spike was conducted in Yanco, Milly-la-Forêt and Sabin in 2018 and Heading date in Milly-la-Forêt and Sabin in 2018.

### QTL mapping of SPS

The 4-parent CSIRO MAGIC population had previously been genotyped using both the Infinium 9k and 90k SNP chips as well multiple DArTs and microsatellites for key genes^32,33^ and the 8-parent NIAB MAGIC population using the Infinium 90k SNP chip^28,34^. QTL analyses were conducted using whole genome average interval mapping using MPWGAIM^33^. For the NIAB MAGIC population a genetic map adapted from Gardner et al., 2016^34^ was used. For the CSIRO MAGIC population a previously published map was used^33^. Skimmed genetic maps containing only unique mapping locations were used to calculate MAGIC RIL haplotypes and founder probabilities, and which were then used for QTL mapping using MPWGAIM. Assigned candidate genes are based on previously published allelic information^35^.

### Development of HIF lines and QTL fine mapping

Following the process described by Barrero et al., 2015^31^, four founders of heterogeneous inbred families (HIFs) were identified from the F_7_ MAGIC RILs through genotyping of the 7AL SPS QTL region using a selection of KASP markers designed based on SNPs from the Infinium iSelect 90K array ^63^. After selfing of each (heterozygous) HIF founder, homozygous progeny with contrasting SPS QTL genotypes were selected and self-pollinated to generate sets of contrasting NILs. Plants from four families were then phenotyped across six field trials as described above.

Genotyping of the F_12_ lines with the Infinium iSelect 90K array^63^ was used to capture any additional recombination in the SPS QTL region and to assign founder genotypes. All NILs were checked and found to contain <1% residual heterozygosity.

### 7AL and 7BL SPS QTL Candidate gene identification

Candidate genes and their orthologues were initially identified in 2016 based on proprietary gene models. Subsequently the regions have been reannotated with the IWGSCv1.1 gene models as shown in Figure 2. Wheat homoeologues and rice orthologues were identified using Ensembl version 51. Orthologue function was obtained through literature search and from funRiceGenes^64^.

### 7AL and 7BL SPS QTL Candidate gene sequencing

Publicly available *WAPO-A1* and *WAPO-B1* sequences were obtained from the reference wheat line Chinese Spring^41^ as well as from Brompton, Robigus and Cadenza (parent of Xi-19)^43^. Additionally, full length sequences for *WAPO-A1* including 5kb upstream promoter sequences in Yitpi, Baxter, Chara and Westonia were obtained via cloning and Sanger Sequencing. Exome sequences for *WAPO1* homoeologues in Alchemy, Claire, Hereward, Rialto, Soissons and Xi-19 were obtained via Exome Capture sequencing.

Based on the Chinese Spring reference sequence primers were designed and *WAPO-A1, B1* and *D1* sequences were cloned from Chinese Spring, Chara, Baxter, Westonia and Yitpi using a Zero Blunt TOPO Cloning kit (Invitrogen). Subsequently the 5kb promoter sequences for *WAPO-A1* were cloned from overlapping PCRs using the same methodology. Clones were selected on LB + Kanamycin medium and sent for Sanger sequencing after two nights of incubation at 37°C. Sanger Sequencing was performed by LGC (UK). Sequences are available under GenBank accession numbers ON210994 to ON210998. PCR conditions and primers are available in Supplementary Table S6.

Wheat Exome Capture sequencing was conducted using a set of internal exome probes based on the IWGSC RefSeqv1 annotations (pre-release) including CDS + UTR sequences. Probe design and balancing was carried out by Roche. Genomic DNA was extracted using a Qiagen DNeasy Mini Kit (Qiagen, US). Library preparation, sequence capture and sequencing were performed at Fasteris. Sequencing was performed on an Illumina HiSeq3000 to obtain an average of 8.2Gb of raw sequence data (2×150bp paired end reads) per sample. Reads for the eight NIAB founders were mapped to the repeat-masked Chinese Spring reference genome (IWGSCv1.0) including mitochondrial (EMBL accession: AP008982) and chloroplast (EMBL accession: AB042240) genomes with BWA^65^ (Li, 2013). This includes an additional remapping step to fine-map InDels. Variant calling and filtering were conducted using GATK^66^. Data was stored and accessed through the BiofacetSNP sequence variant data management system.

### Candidate gene expression analysis

Chinese Spring developmental time course gene expression data from Choulet et al., 2014^67^ for corresponding TGACv1 gene models was accessed on wheat-expression.com^45^ in March, 2017. As *WAPO-A1* was not annotated as part of the IWGSC_CSSv2.2^68^ and TGACv1 gene models^69^ available B and D genome models were used as a query instead.

### Gene expression of *WAPO-A1* in the field

Three replicates of whole spike samples were collected from tiller dissections of the 2017 NIAB MAGIC Nursery at growth stage GS32^44^ for the MAGIC founders Alchemy, Brompton, Claire, Hereward, Rialto, Robigus and Xi-19. At the collection date Soissons had advanced to GS34. Following dissection, spikes were immediately frozen in liquid nitrogen. For *WAPO-A1*, betaine solution was added at a final concentration of 1M to overcome the amplicons high GC content. To confirm specificity of qRT-PCR reactions the melt curves for each reaction were checked for the presence of only a single peak. Specificity of the assays was confirmed against genomic nullitetrasomic DNA obtained from Seedstor.ac.uk (WPGS1289-PG-1, WPGS1296-PG-1, WPGS1301-PG-1). Expression levels of W*APO-A1* were calculated relative to the expression of the housekeeping genes *TaRP15*^46^and *Ta2291*^47^. Primers sequences are in Supplementary Table S7.

### *WAPO-A1* Transformation experiments

A 7,973 bp gene cassette containing the entire Westonia *WAPO-A1* sequence including 5.4 kb of the native promoter was synthetically design at GeneArt (Thermo Fischer Scientific, US) and received in the standard cloning vector pUC57. This cassette was subcloned into the binary vector VecBarII (using the XbaI and HindIII sites), which has been extensively used at CSIRO for wheat transformation^70^. CSIRO’s Transformation Unit performed the transformation of the Yitpi cultivar or Fielder, using Agrobacterium^71^. Yitpi was chosen over Chara for the complementation experiment because of previous work done at CSIRO showing higher transformation rates in Yitpi (19-27% for Yitpi vs. 5-9% for Chara, data not shown).

Thirteen Yitpi and five Fielder primary transformants were generated and grown in pots in the glasshouse. To assess the copy number in each primary transgenic, 25 T_1_ seeds from each T_0_ plant were germinated in petri dishes and a Basta-resistance ‘leaf assay’ was used to determine transgene segregation^72^. After a week, the resistance to Basta was scored to determine segregation ratios (data not shown). Based on the segregation ratios six T_0_ Yitpi and three T_0_ Fielder transformants were selected for further work. From each of them eight T_1_ positive lines and two null-segregants were transplanted to soil and grown in glasshouse conditions. For each T_1_ plant, about 25 T_2_ seeds were used in leaf-tipping assays to identify homozygote lines. A T_2_ homozygous and a null-segregant lines from each family were chosen for final analysis. Ten plants of each were transplanted to pots in the glasshouse and they were phenotyped at maturity. Samples were also collected for gene expression analysis in developing spikes.

Gene expression of WAPO-A1 in transgenic and control lines was analyzed by RT-qPCR in developing spikes at GS32. Expression levels were calculated relative to the housekeeping gene TaActin1 (CJ961169). Primer sequences are in Supplementary Table S7.

All transgenic work was conducted at CSIRO in Canberra (ACT, Australia) in certified Physical Containment Level 2 (PC2) facilities, including laboratories and glasshouses, following the standards set by the Australian Office of the Gene Technology Regulator (https://www.ogtr.gov.au/).

## Acknowledgements

CSIRO acknowledges the contributions of Dr Megan N. Hemming, Dr Alex Whan, Dr Richard A. James, Dr Sally Walford and Mr Andrew Spriggs for their support at the early stages of this project. Thanks to Ms Tracy Willis, Mr Jackson Tahapehi, Mr Alexandre Boyer, Dr Aderajew Haddis, Ms Alison Fattore, Ms Bec Maher, Mr David Lewis, Mr Max Bloomfield, Mr Mick Weiss, Mr Patrick Moody and Mr Todd Collins for the management of the field trials at CSIRO. We also thank Ms Terese Richardson and Ms Dhara Bhatt for their work at the CSIRO Wheat Transformation Unit. BASF thanks Xi Wang and Ruvini Ariyadasa for contributions to data analysis early in the work and Greta De Both, Stephanie Thepot, and David Bonnett for management of the field trials at BASF. This project was supported by BASF. Lukas Wittern was funded by BBSRC iCASE grant BB/K011790/1 awarded to Alex Webb (University of Cambridge), Matthew Hannah (BASF) and Andy Greenland (NIAB). Gareth Steed was funded by BBSRC iCASE grant BB/M015416/1 awarded to Alex Webb and Matthew Hannah.

## Competing interests

Patent application ^73^WO2019/138083 A1 has previously been submitted based on some of these results for which Mark Davey, Colin Cavanagh, Matthew Hannah, Lukas Wittern, Keith Gardner, William Bovill, Jose Barrero and Alex Webb are among inventors. John Jacobs, Mark Davey, Matthew Hannah, Claus Frohberg, Ralf-Christian Schmidt, Colin Cavanagh and Antje Rohde are employees of BASF. Andy Greenland, Gareth Steed, Steve Swain and Trijntje Hughes declare no competing interest.

## Data availability

The datasets generated are available for academic partners for non-commercial purposes upon request sent to the corresponding author, provided that bilateral terms-of-use agreements can be concluded.

## Author contributions statements

The 4-parent CSIRO MAGIC QTL and HIF work was led by WDB and KLV. JMB and TH did the transgenic analysis. Input on design, analysis and interpretation of results was provided by AR, CC, CF, JJ, MD, MH, RCS and SS. The 8-parent NIAB MAGIC QTL work was conducted by LMW, with gene-expression analysis by LMW and GS. Input on design, analysis and interpretation of results was provided by AARW, KG, AG and MH. LMW and JMB wrote the manuscript. All authors reviewed the manuscript.

## Notes

### Competing Interest Statement

Patent application WO2019/138083 A1 has previously been submitted based on some of these results for which Mark Davey, Colin Cavanagh, Matthew Hannah, Lukas Wittern, Keith Gardner, William Bovill, Jose Barrero and Alex Webb are among inventors. John Jacobs, Mark Davey, Matthew Hannah, Claus Frohberg, Ralf-Christian Schmidt, Colin Cavanagh and Antje Rohde are employees of BASF. Andy Greenland, Gareth Steed, Steve Swain and Trijntje Hughes declare no competing interest.

### Summary of Updates

Body text revised; supplemental tables updated; author list corrected.

