## Supplementary Figures for "Overexpression of the *WAPO-A1* gene increases the number of spikelets per spike in bread wheat"

### Appendix

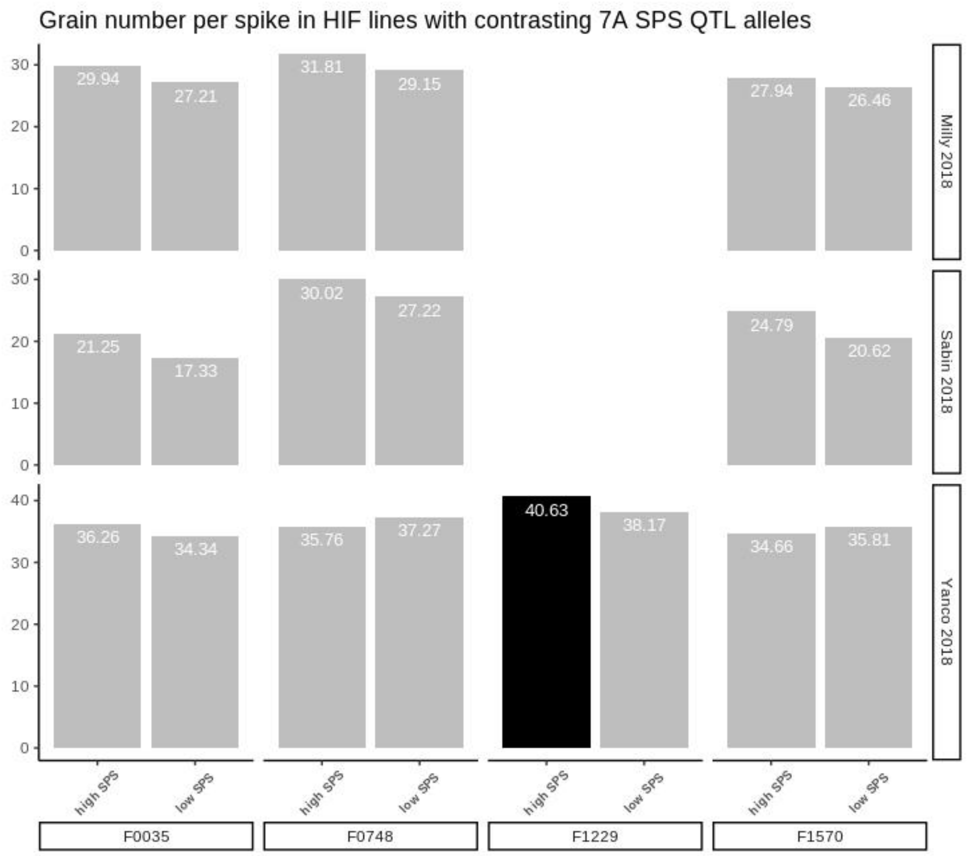


**Supplementary Figure S1: Validation of phenotypes for 7AL SPS QTL using HIF families.** A) Summary of differences in grain number per spike across NIL pairs for the 7AL SPS QTL. Black fill indicates a significant increase ($p < 0.05$) between NIL pairs.

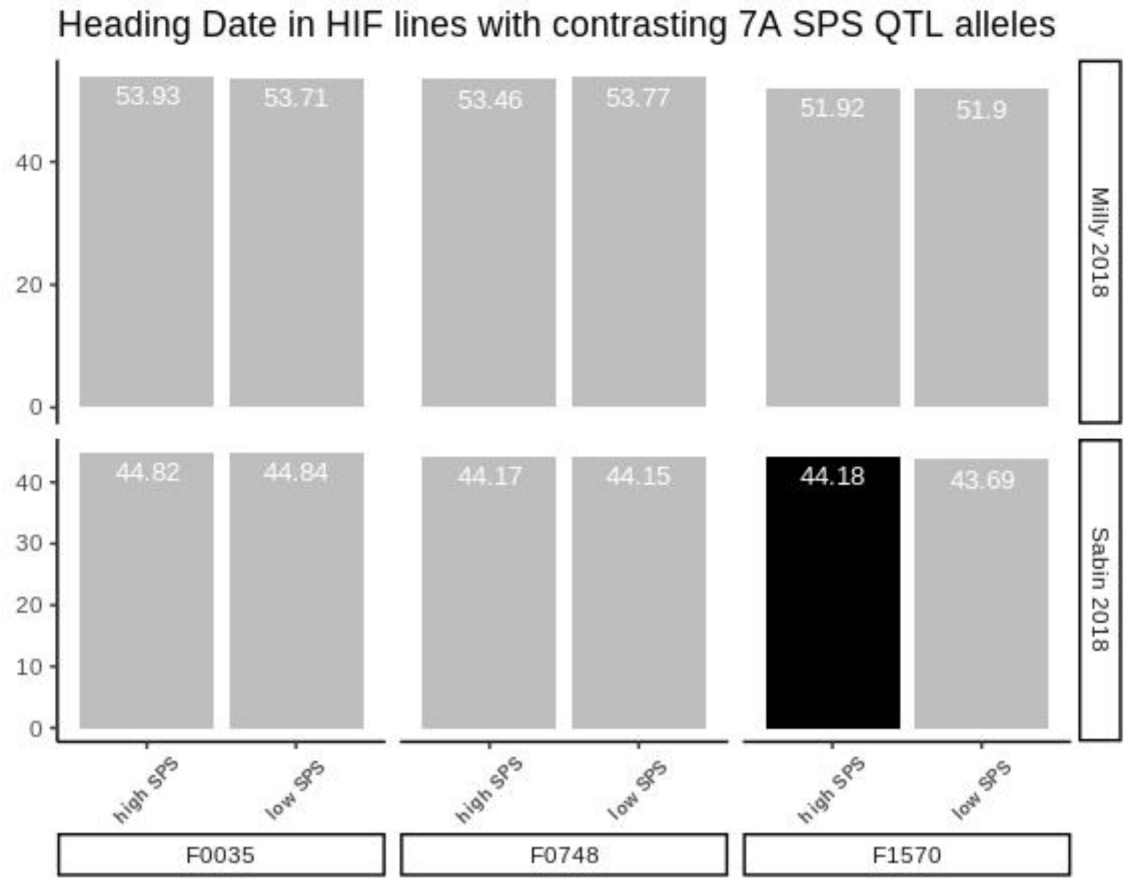


**Supplementary Figure S2: Validation of phenotypes for 7AL SPS QTL using HIF families.** A) Summary of differences in heading date across NIL pairs for the 7AL SPS QTL. Black fill indicates a significant increase (p<0.05) between NIL pairs.

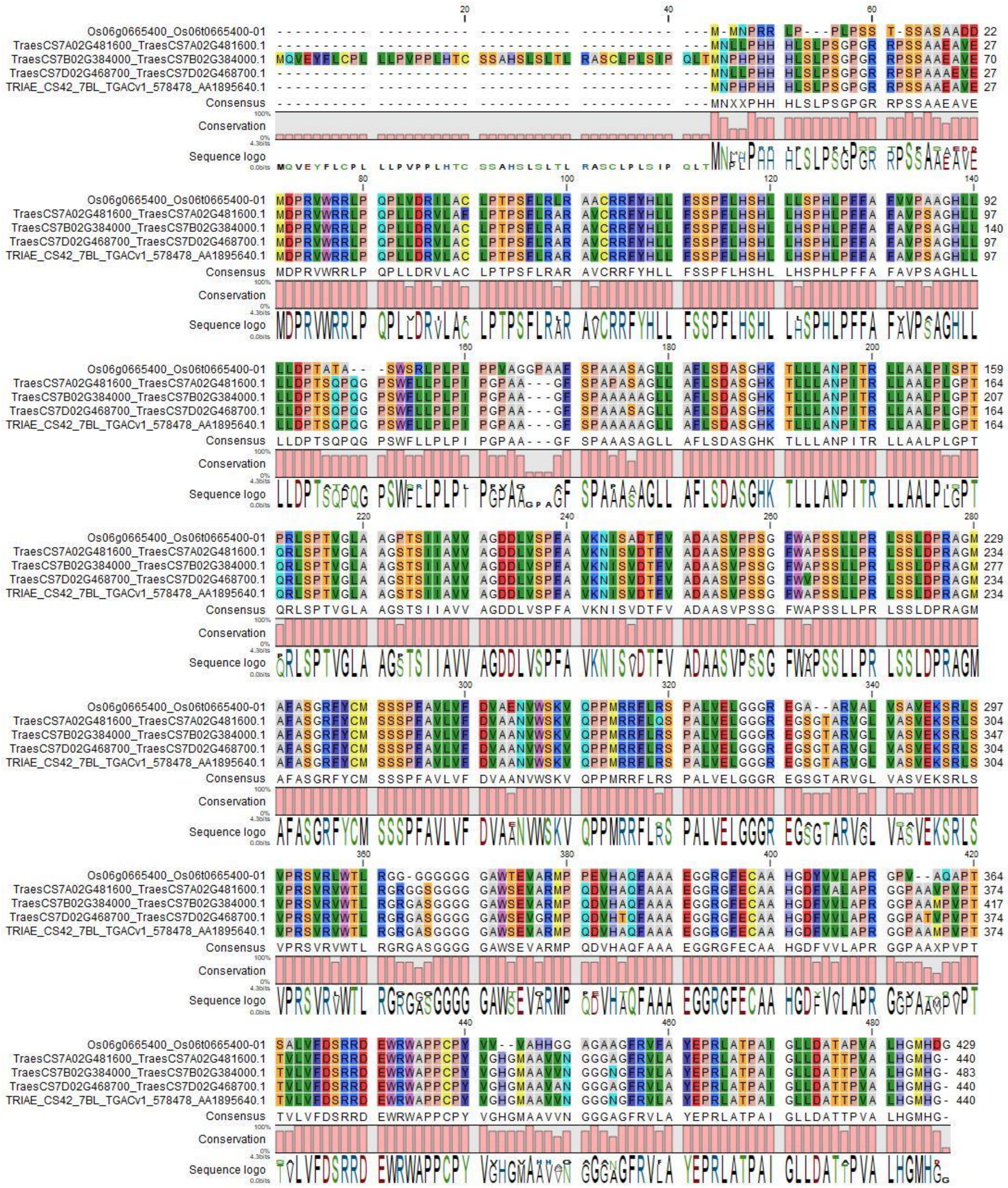


**Supplementary Figure S3: Alignment of WAPO1 homoeologues and OsAPO1 (Os06g0665400).** MCoffee alignment visualised in CLC Genomics Workbench. Amino acids coloured based on RasMol colour scheme. Amino acid % identity conservation between *WAPO-A1* and *OsAPO1* is 80%. The alternative gene model for *WAPO-B1* is *TRIAE_CS42_7BL_TGACv1_578478_AA1895640.1.*

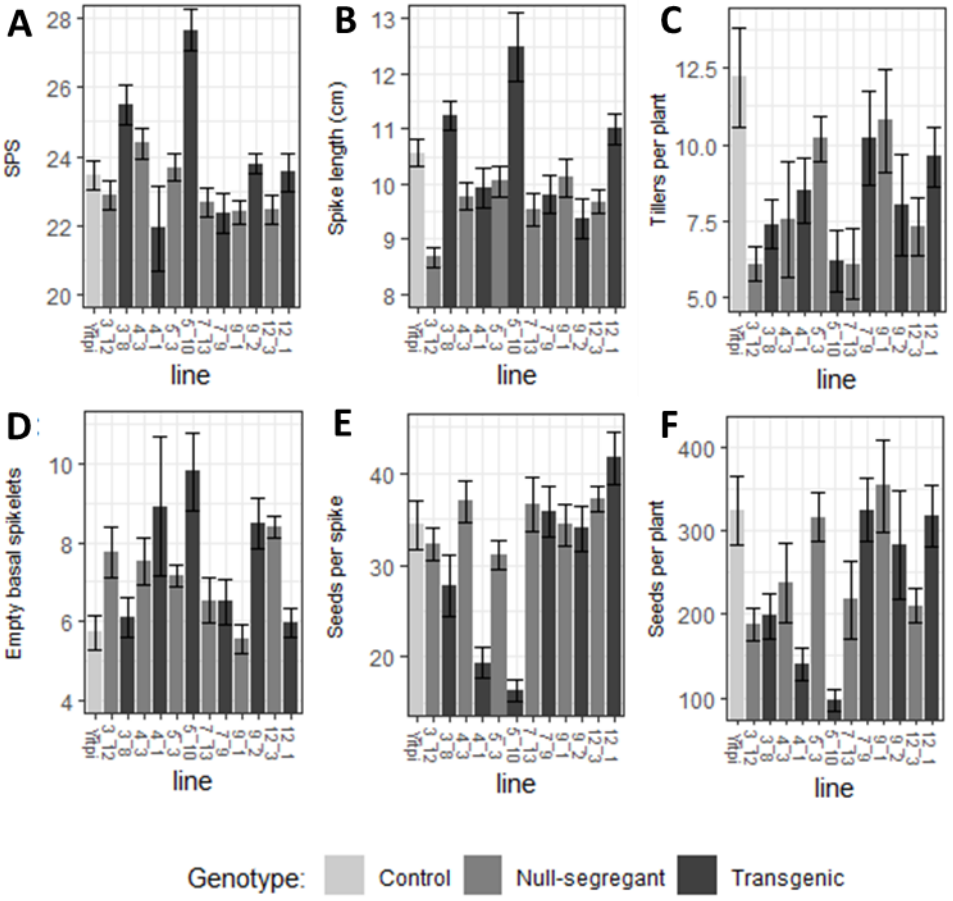


**Supplementary Figure S4: Phenotypic characterisation of Yitpi transgenic, null-segregates and control spikes and plants.** A) SPS number. B) Spike length. C) Tiller number per plant. D) Empty basal spikelets. E) Seeds per spike. F) Number of seeds per plant. Means with SEM are represented. For spike traits, 50 spikes (5 spikes x 10 plants) of each genotype were used in the calculations. Note variation in range for Y axes.

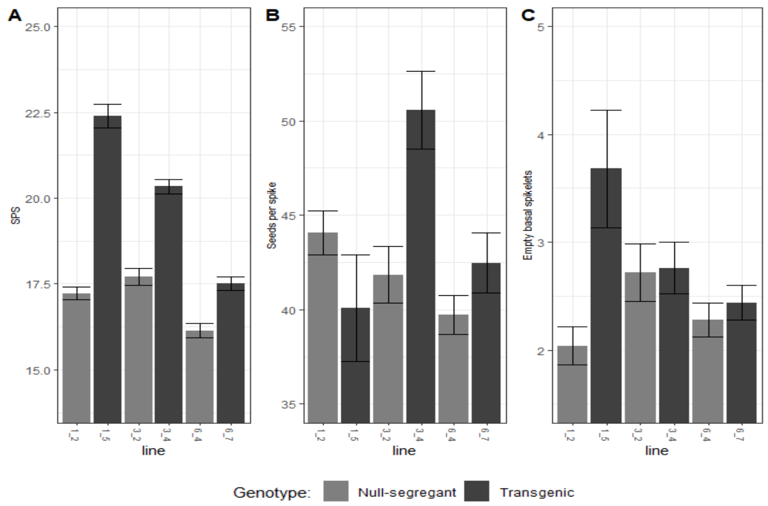


**Supplementary Figure S5: Spike phenotype of transgenic Fielder plants and null-segregants.** A) SPS. B) Seeds per spike. C) Empty basal spikelets. Means with SEM are shown. 50 spikes (5 spikes x 10 plants) of each genotype were used in the calculations.

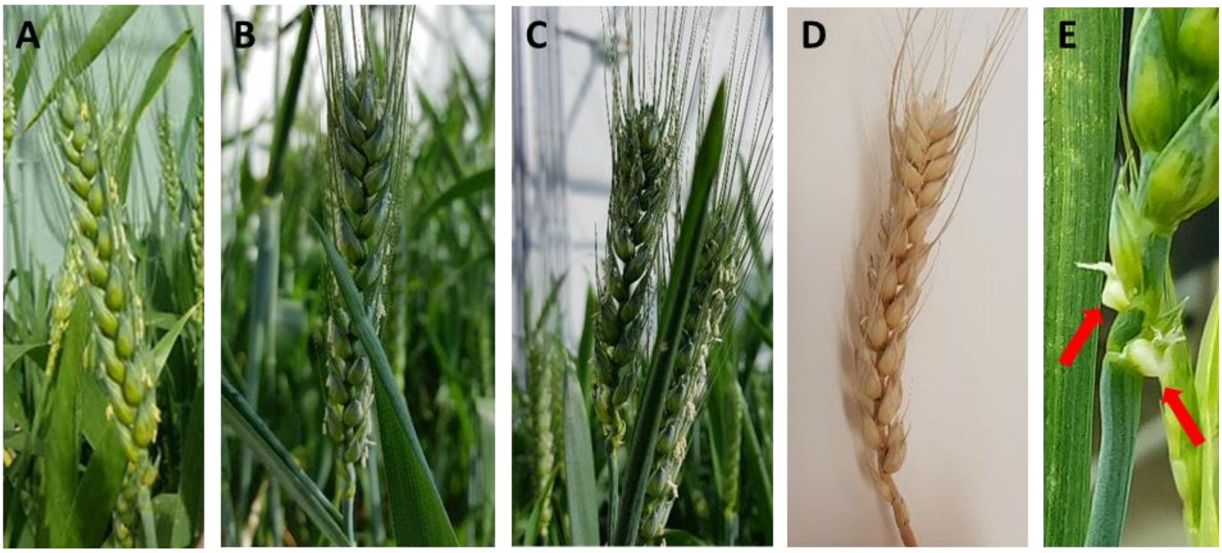


**Supplementary Figure S6: Examples of compact spike phenotype and of basal floret defects in Fielder transgenics.** A) Fielder control spike. B-D) Examples of compact heads in over-expression lines. E) Details of homeotic floral defect in basal florets.

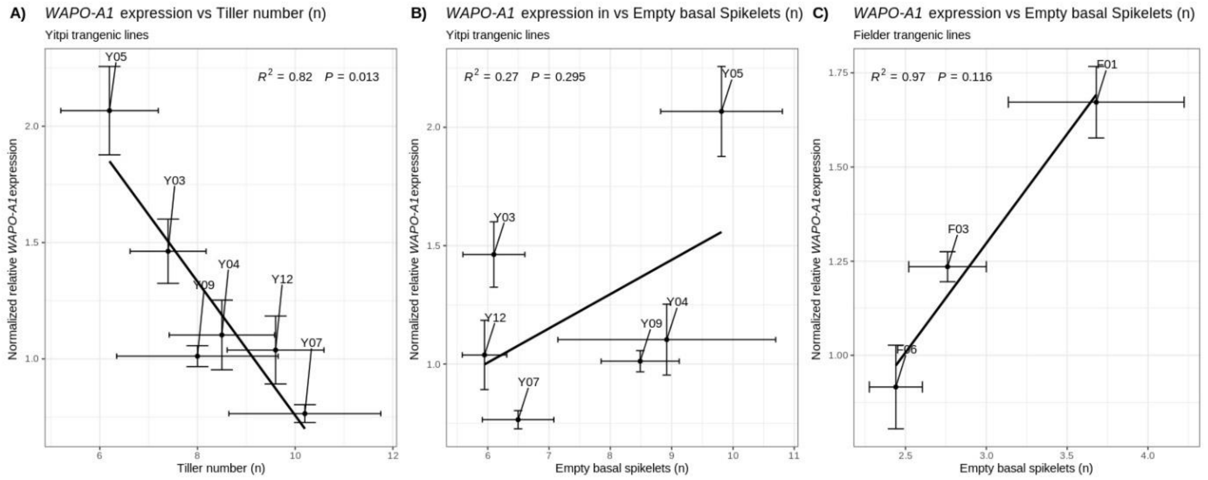


**Supplementary Figure S7: Effect of WAPO-A1 expression on Tiller number and Empty basal spikelets.** Gene expression was assayed in developing spikes dissected at GS32. A) Regression of normalized expression of *WAPO-A1* on Tiller number measured in Yitpi transgenic lines. B) Regression of normalized expression of *WAPO-A1* on Empty Basal Spikelets measured in Yitpi transgenic lines. C) Regression of normalized expression of *WAPO-A1* on Empty Basal Spikelets measured in Fielder transgenic lines. Means and SEM of three biological replicates are shown.

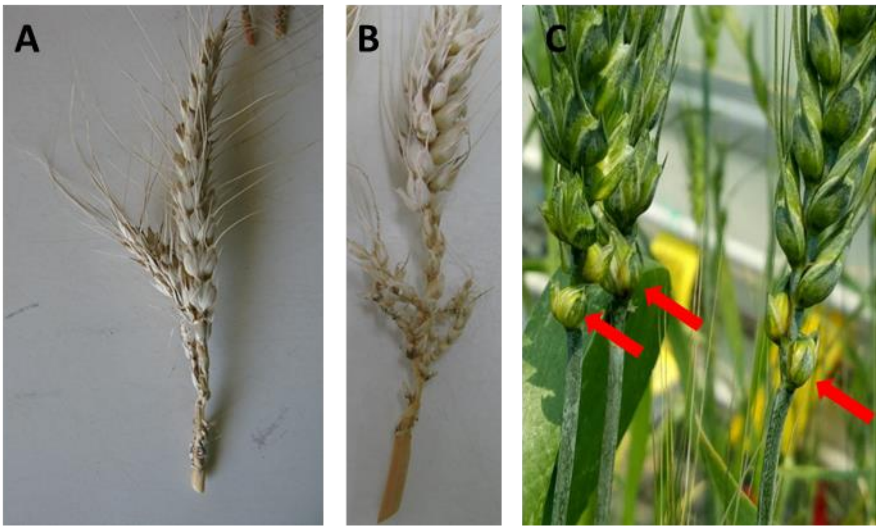


**Supplementary Figure S8: Examples of branching spikes and floral defects in transgenic line 5-10.**
A-B) Extreme branching phenotype. C) Details of homeotic floral defect in basal florets
